# Parasitized or non-parasitized, why? A study of factors influencing tick burden in roe deer neonates

**DOI:** 10.1101/2022.01.11.475805

**Authors:** Léa Bariod, Sonia Saïd, Clément Calenge, Stéphane Chabot, Vincent Badeau, Gilles Bourgoin

**Affiliations:** Université de Lyon, VetAgro Sup - Campus Vétérinaire de Lyon, Laboratoire de Parasitologie Vétérinaire, Marcy-L’Etoile, France; Université de Lyon, Université Lyon 1, CNRS, UMR 5558, Laboratoire de Biométrie et Biologie Évolutive, Villeurbanne, France; Office Français de la Biodiversité, Direction de la Recherche et de l’Appui Scientifique, Birieux, France; Office Français de la Biodiversité, Direction de la Recherche et de l’Appui Scientifique, Le Perray en Yvelines, France; Institut National de Recherche pour l’Agriculture, l’Alimentation et l’Environnement, Champenoux, France

**Keywords:** *Capreolus capreolus*, newborns, *Ixodes ricinus*, age, climatic variability, habitat, population density

## Abstract

*Ixodes ricinus*, the most common species of tick in Europe, is known to transmit major pathogens to animals and humans such as *Babesia* spp. or *Borrelia* spp.. Its abundance and distribution have been steadily increasing in Europe during recent decades, due to global environmental changes. Indeed, as ticks spend most of their life in the environment, their activity and life cycle are highly dependent on environmental conditions, and therefore on climate or habitat changes. Simultaneously, wild ungulates have expanded their range and increased dramatically in abundance worldwide, in particular roe deer (*Capreolus capreolus*), have allowed tick populations to grow and spread. Currently, tick infestation on newborn wild ungulates is poorly documented. However, newborn ungulates are considered more sensitive to tick bites and pathogen transmission because of their immature immune system. Thus, improving knowledge about the factors influencing tick infestation on newborns is essential to better understand their health risks. This study was conducted at Trois-Fontaines forest, Champagne-Ardenne, France (1992-2018). Based on a long-term monitoring of roe deer fawns, we used a novel Bayesian model of the infestation of fawns to identify which biotic or abiotic factors are likely to modify the level of infestation by ticks of 965 fawns over time. We show that tick burden increased faster during the first days of life of the fawns and became constant when fawns were 5 days old and more, which could be explained by the depletion of questing ticks or the turnover of ticks feeding on fawns. Moreover, the humidity, which favors tick activity, was weakly positively related to the tick burden. Our results demonstrate that tick infestation was highly variable among years, particularly between 2000 and 2009. We hypothesize that this results from a modification of habitat caused by hurricane Lothar.

## Introduction

Ticks are considered to be one of the most important arthropod vector of diseases in the world [1]. They are of major concern as they can transmit several pathogens to their host during their blood meal, such as viruses, bacteria or protozoans [2,3]. They can also have direct negative effects to their host, such as blood loss (known as spoliation effect [4]) or destruction of host tissue by enzymes present in their saliva [5]. Over the past decades, these ectoparasites have become an increasing worldwide health burden, threatening both humans and animals [6]. The abundance and distribution of ticks can strongly vary among areas and have considerably increased in recent decades [7–9]. In order to understand the population dynamic and the geographic distribution of ticks, and to predict the risks of transmission of tick-borne pathogens, it is of major concern to determine the factors driving abundance and spread of these parasites [10].

In Central Europe, *Ixodes ricinus* is the most common hard-tick species. It is of particular concern due to their major role as a vector of many pathogens for both animals and humans, such as *Anaplasma* spp. [11], *Babesia* spp. [12], *Borrelia* spp. [13] and Tick-borne encephalitis virus [14]. This three-host tick species takes one blood meal on their host per life stage (3 life stages: larva, nymph and adult), with blood meals lasting for few days (2-12 days according to their life-stage; [15,16]) before falling in the environment where it moults into the next stage or lay eggs and die [17]. As this tick completes its life cycle in at least 3 years [15], it spends most its life free in the environment. Thereby survival, development and questing activity are highly dependent on the weather conditions and microclimate [18,19].

Among the microclimatic requirements, temperature and humidity are the most important factors affecting the development rate, survival and questing activity of ticks [20]. Some studies have estimated that *Ixodes ricinus* is looking for a host when the temperature is in average between 7°C and 26°C [21–24] and when the relative humidity is at least 80% to avoid desiccation [7]. When these conditions are met, ticks use the “ambush” technique in climbing up vegetation to quest for a passing host [7]. However, during this activity step, ticks loose moisture, which obliges them to return often to the ground to recover the lost water. The suitability of the ground based-vegetation which retains the moisture and provides shadow is highly important for the tick survival, ticks are abundant in the leaf litter and the low strata vegetation of woodlands and forests [7,25]. *I. ricinus* can survive in deciduous forests or in meadows as long as there is sufficient litter on the ground and the microclimate is moist [26]. However, it has been shown that the abundance of these ticks is higher in deciduous forests, particularly those containing oak and beech, and especially when the shrub cover is important [26,27]. Local microclimate and local vegetation have thus important roles for tick survival and behavior in their seasonal questing activity throughout the year [22,28,29]. In general, a bimodal activity pattern with a dominant peak in spring and a minor peak in autumn is observed for *I. ricinus* in Central Europe when climatic conditions are temperate and wet. In addition, as variations of seasonal weather conditions between years occur, yearly variations of the seasonality of tick activity should also occur [24].

The life cycle of *I. ricinus* is also highly dependent on the availability of hosts, because of their questing behavior (i.e., the “ambush” technique; [7]). This ectoparasite species is a ubiquitous tick in Europe, which can parasite a wide range of animal species, including mammals, reptiles and birds [30]. Larvae and nymphs of *I. ricinus* are commonly found on small animals such as rodents [31], although nymphs can also be seen on larger hosts such as sheep [32]. Large vertebrates are the main suitable hosts for blood meals and mating of the adult ticks [26]. The infestation consequences on host will be greater especially if these parasites are found in large quantities [4].

Hosts can develop an immune response against ticks to decrease their fixation time and engorgement, and hence, the spoliation effect and the risk of pathogen transmission [33,34]. Resistance to tick infestation implicates both acquired and innate immunity [35]. However, ticks can circumvent host defenses with active components secreted in their saliva which can induce host immunosuppression and facilitate acquisition of blood meals for ticks and the transmission of the tick-borne pathogens [36]. This is why newborns can be considered more sensitive to tick bites. Indeed, they have a naïve and immature immune system even if they are not totally vulnerable thanks to the transmission of acquired immunity by the colostrum of the mother [37,38]. In addition, failure in the transfer of humoral immunity from mothers can happen and increase susceptibility of newborns to tick bites [39,40].

The cervids, and particularly the roe deer (*Capreolus capreolus*), are considered as key hosts for the population dynamic of *I. ricinus* [26]. During the last decades, their population have been increasing in the Northern Hemisphere [41,42], which, in turn, has led to the increase and geographical spread of tick populations [10,26,43–45]. Tick activity is the most important in spring and the eggs laying by female ticks takes place at the same time as the birth period of the fawns, *i*.*e*. around the month of May [24,46]. Furthermore, fawns of roe deer have a hider behavior (i.e., they hide in the vegetation cover to limit the probability of detection by a predator; [47]) and have therefore a preference for dense vegetation. Their naïve immune system make also them an easy target for questing ticks even if their movements are limited during the first days of life [48,49]. To our knowledge, no study has yet investigated the factors influencing the level of infestation by ticks on roe deer neonates.

Based on previous knowledge on ticks, we predicted a higher tick burden (H1) during warm and wet spring conditions as it favors tick activity. We also predicted (H2) an influence of the habitat on tick infestation, with more parasitized roe deer fawns living in habitats with important shrub cover and ground-based vegetation which retains humidity; and (H3) an influence of the density of roe deer on tick burden, with the highest values of infestation by ticks during the years when roe deer density, calculated the winter preceding the catches of fawns, is the highest. Indeed, as intensity of tick burden on deer increases with high host populations densities, we can assume that ticks will be more abundant in the environment and parasitize fawns more [10,50].

Based on a long-term monitoring of the roe deer neonates from 1992 to 2018 in the population of Trois-Fontaines in France, we studied the factors influencing the tick burden of newborn roe deer fawns. This long-term monitoring of the infestation of fawns during their first days of life by the tick *Ixodes ricinus* allowed us to study the effects of factors linked to the host (individual and populational factors) on tick risks, but also of variations in Spring weather conditions (temperature, humidity, extreme events) and habitat characteristics. We used a novel Bayesian model of the infestation of fawns by ticks to identify which biotic or abiotic factors are likely to level of tick infestation of fawns over time.

## Materials and methods

### Ethics statement

In accordance with European and French laws, roe deer captures have been carried out minimizing animal stress and handling time (limited to 10 minutes for a fawn), and ensuring animal welfare, as defined in the guidelines for the ethical use of animals in research. All methods were approved by the authorities (French Ministry of Environment). These experiments were performed in accordance with the conditions detailed in the specific accreditation delivered to the Office National de la Chasse by the Préfecture de Paris. Animal captures and experimental procedures were in line with the French Environmental Code (Art. R421-15 to 421–31 and R422-92 to 422-94-1) and duly approved by legislation from the Prefecture of Paris (agreement n°2009–014, n°. 2013–118, n°2019-02-19-003).

## Data collection

### Study area

This study was carried out in the Territoire d’Étude et d’Expérimentation (TEE) of Trois-Fontaines, located in north-eastern France (48°38 ‘ N, 4°54 ‘ E) (Fig 1). This is an enclosed forest of 1360 ha with an overstory dominated by oak (*Quercus* spp.) and beech (*Fagus sylvatica*), while the coppice is dominated by hornbeam (*Carpinus betulus*) [51]. The climate is continental, characterized by cold winters (mean daily temperature in January-February was 3.2°C between 1992 and 2018, data from Météo France) and hot but not dry summers (mean daily temperature in July is 19.2°C and total rainfall in July and August is 137.63 mm between 1992 and 2018, data from Météo France).

**Fig 1.**
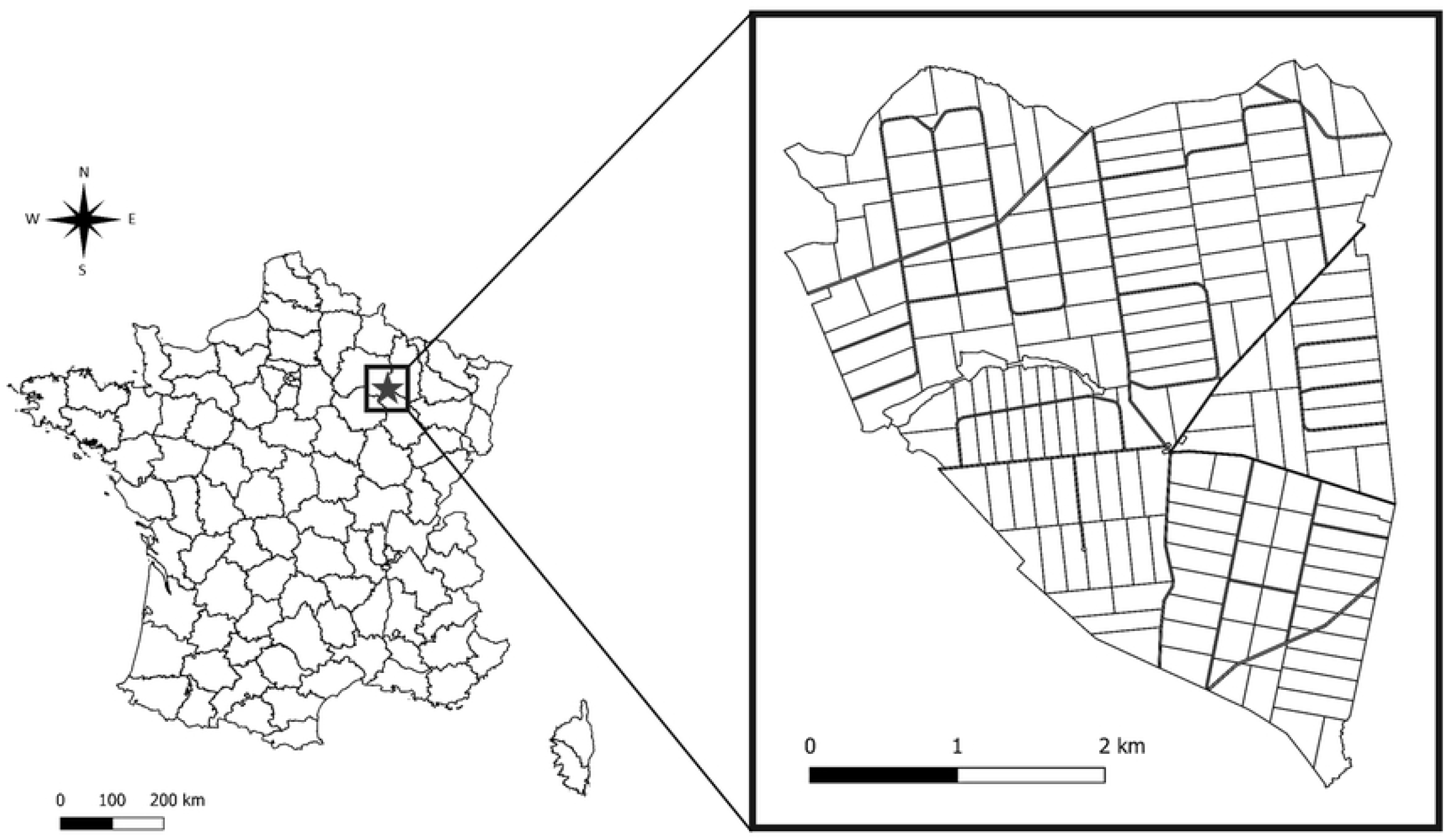
Geographic location in France (on the left) of the study site composed of 172 forest plots (on the right).

### Fawn data and tick sampling

Search of fawns were organized in the TEE of Trois-Fontaines by field workers and volunteers (hunters, foresters, etc.) every year during the fawning period (April to June) [46]. Fawns detected hiding in a bedding site during their first days of life were captured. Then, we identified each fawn with ear-tags and collected information on sex, age (in days) and body mass (in kg). Umbilicus characteristics and behavior of fawns during the capture were used to determine their age [52].

We assessed the level of tick infestation by visually and manually inspecting the neck and head of each captured fawn. Based on the number of detected ticks, we attributed to each fawn the following infestation classes: (i) class 1 = < 10 ticks; (ii) class 2 = 10 - 20 ticks; (iii) class 3 = > 20 ticks. A previous study of the ticks in Trois-Fontaines showed that *Ixodes ricinus* was the only tick species observed on roe deer (G. Bourgoin, *unpublished data*).

Based on their measured body mass, we calculated the body surface area (bsa) of each fawn using the Meeh’s formula [53]:

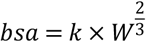

with k: Meeh constant (k=10); W: body mass (in kg).

### Data on the population of roe deer and the environment

The roe deer population in Trois-Fontaines has been intensively monitored for 40 years using capture-mark-recapture methods [54]. Driving nets are used each year in December-March to catch the individuals (10-12 days of capture per year). Using CMR modelling, the number of roe deer older than one year was thus estimated every year in March, which we could use to derive a value of density every year (Fig 2) [55].

**Fig 2.**
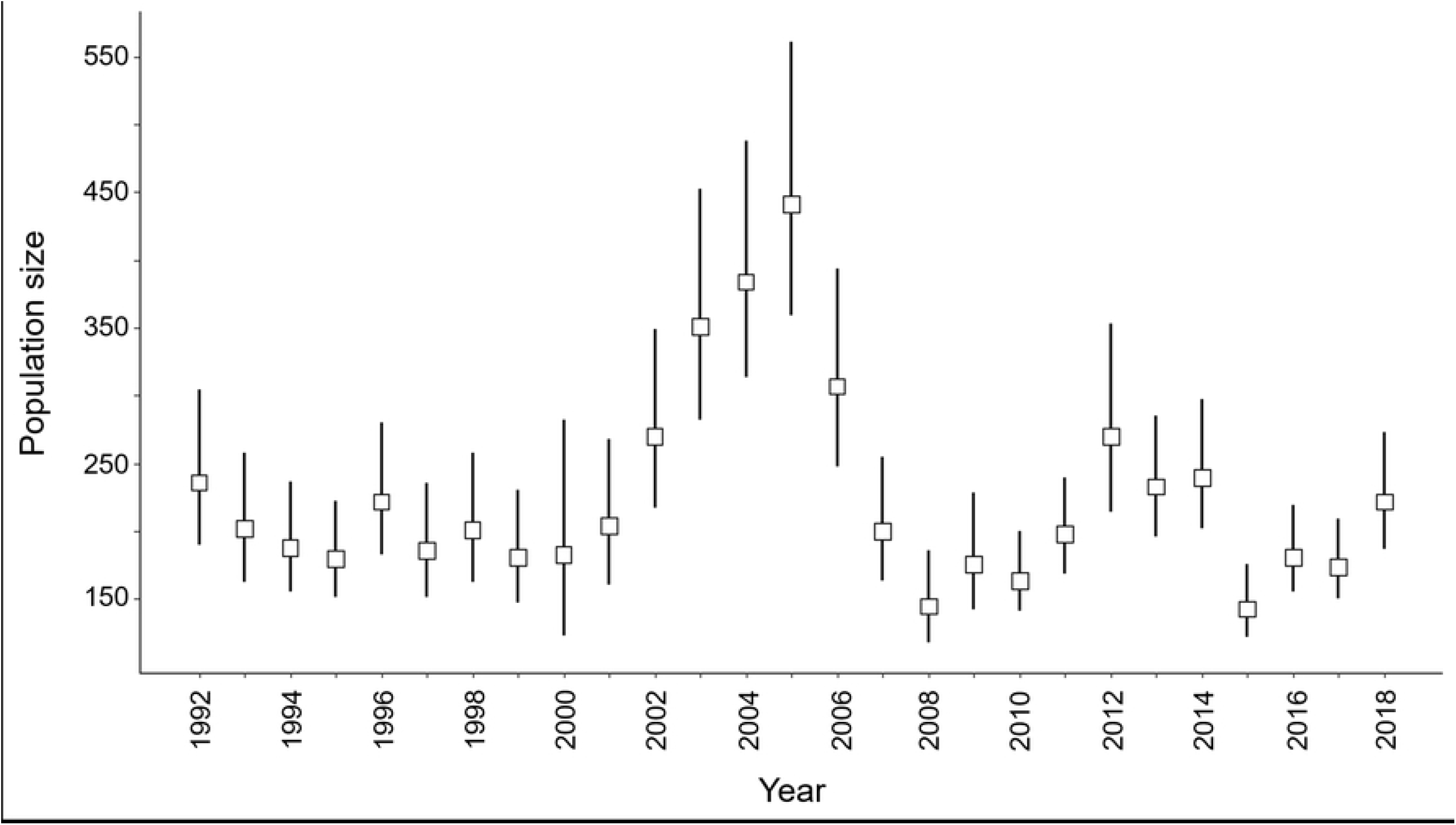
Yearly estimates (with 95% confidence intervals) of roe deer population density in March individuals > 1 year) at TEE of Trois-Fontaines (France) estimated by capture-mark-recapture models (see [54] for further details) for the period 1992–2018.

Weather data of this study was obtained thanks to the SAFRAN system. This is a mesoscale atmospheric analysis system that reconstructs surface atmospheric variables at the hourly timescale on a regular 8 km grid using ground observations and large-scale general circulation model products (Météo France, [56]). This reanalysis is described in Durand et al. [57,58] and several validations have been performed [56,59]. In this study, we used the mean values of the temperature (T, in °C) and relative humidity (RH, in %) of the capture day and 4 days before this day. Indeed, nymph and larvae of ticks take a blood meal on their host during in average 3 days, and adult in 7 days, before to detach and fall off [16]. We therefore considered a mean duration of blood meal of 5 days, explaining why we took the weather conditions into account during this time.

Our study site contains 172 forest plots measuring in average 7.95 ha, delineated by forest trails. In each forest plot during autumn, the dominant species and its cover proportion were determined for the following years: yearly from 1996 to 2005, in 2009 and in 2012 [51,58]. We used these data to assess the habitat effect on tick infestation. When habitat data was not available for a year: (1) we used habitat data from the closest year (i.e., +/-2 years max); (2) data after 2014 were not considered, because we did not collect any data on habitat structure after 2012, and we considered that habitat structure had changed too much after 2 years to be ignored in a study of habitat effect on tick infestation.

Between 1996 and 1999, forest managers have made openings in the habitat (i.e., pasture area) to improve roe deer habitat. Then, in late December 1999, hurricane Lothar, apparently the strongest hurricane in the region for at least 1000 years, hit the Trois-Fontaines site, disturbing the habitat [59,60]. This event created several openings and modified the microclimatic conditions of the environment.

### Model of the infestation process

We designed a Bayesian model describing the infestation process of the roe deer fawns by ticks during the first 8 days of their life, to test the effect of environmental variables on the infestation rate. All the code and data used for this modelling approach are available in a R package named `tickTF` [63], (Digital Object Identifier: 10.5281/zenodo.5764798), available on Github at the URL: https://github.com/ClementCalenge/tickTF. It can be installed in R with the package devtools [64], using the function `devtools::install_github(“ClementCalenge/tickTF”, ref=“main”)`. The package tickTF includes a vignette describing how the user can reproduce easily the model fit (available with the command `vignette(“tickTF”)`, once the package has been installed). This vignette is also available as the supplementary material of this paper, and describes the complete mathematical development of the model, the implementation of the model using the R package `NIMBLE` [65], the complementary analyses and model checks.

In this section, we give a short description of the model, and we refer the reader to the supplementary material for a more complete development.

We modelled the infestation process by a Poisson process (e.g. [66]). In other words, the number *N*_*j*_ of ticks present on a given fawn *j* is supposed to follow a Poisson distribution with mean *Λ*_*j*_:

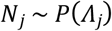

The expected number of ticks on fawn *j* is supposed to be the result of a process of accumulation of ticks, i.e. is equal to the integration over time *t* of an instantaneous risk function *λ*_*j*_*(t)*:

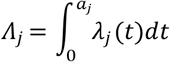

where *a*_*j*_ is the age of fawn *j* at the time of capture. We modelled this instantaneous risk of infection of a fawn having *t* days old with:

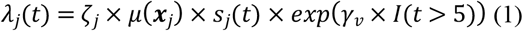

The instantaneous rate of infestation of a given fawn is therefore the product of an instantaneous rate of infestation per unit of body size area *μ*(**x**_j_) -- which itself is supposed to depend on environmental variables characterizing the capture **x**_j_ (see below) --, of the body size *s*_j_(*t*) of the fawn *j* at times *t*, and of a random individual effect *ζ*_*j*_. These random effects follow a gamma distribution G(1/*φ, φ*), with *φ* a dispersion parameter to be estimated, and account for the fact that different fawns have different sensibilities to ticks.

As we said before, we considered a mean duration of blood meal of 5 days. Then, after 5 days, the ticks that infested the fawn on its first day of life start to detach themselves and fall off, which is expected to lead to a decrease in the observed infestation rate, as the number of ticks infesting a fawn may be partly compensated by the number of ticks quitting the animal. Thus, the instantaneous rate of infection is supposed to be different for fawns aged up to 5 days old and for older fawns (this rate is multiplied by exp(γ_v_) after that) – in this equation, *I*(*t*>5) takes the value 1 when the fawn is older than 5 days old and 0 otherwise.

To fit this model, we had to develop a submodel for the instantaneous rate of infestation per unit of body size area *μ*(**x**_j_) – which is independent of age – and another submodel for the changes of the body surface area *s*_*j*_(*t*) of fawn j with time – which varies with age.

For the latter, we could use the data collected on captured fawns to model the relationship between the age at capture *a*_*j*_ and the body surface area at capture *<u>s</u>*_*<u>j</u>*_(*a*_*j*_). Our data suggested that the following quadratic regression model was reasonable to describe the growth of the roe deer fawn during its first 8 days of life (see supplementary material):

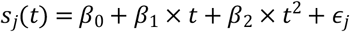

where <u>ε</u>_j_ is a gaussian residual with zero mean and standard deviation σ_s_.

Moreover, we proposed the following loglinear submodel for the instantaneous rate of infestation per unit of body size area:

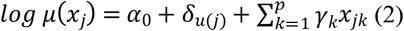

where *α*_*0*_ is the intercept, *δ*_*u*(j)_ is a gaussian random effect characterizing the year *u*(*j*) of capture of fawn *j*, and γ_k_ are coefficients characterizing the effect of environmental variables *x*_*jk*_. Note that preliminary versions of our model revealed that the variance of the year random effects *δ*_*u*(j)_ varied a lot across periods. We distinguished three periods based on the dynamics of the forest structure: (i) period 1 corresponded to the period before the Lothar hurricane (between 1992–1999), (ii) period 2 corresponded to the 10 years following this hurricane (between 2000 and 2009), (iii) period 3 corresponded to the later years (2010 to 2018). We estimated one variance parameter 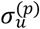 of the year random effects for each period *p*. During the step of model building, we tested the effect of several variables *x*_*jk*_ on the infestation rate: (i) humidity, (ii) temperature, (iii) roe deer density and (iv) habitat type. We kept in the model only the most influential variables.

One difficulty with this model is that we did not know the exact number *<u>N</u>*_*j*_ of ticks on the fawn *<u>j</u>* but rather the tick-burden class. However, given *Λ*_*<u>j</u>*_ we could calculate the probability of all the possible values of *<u>N</u>*_*<u>j</u>*_ within a given class (since *N*_*i*_ follows a Poisson distribution), and therefore, we could calculate the probability of each burden class by summing the probabilities of all values of *<u>N</u>*_*j*_ in this class (see supplementary material for more details).

We estimated the posterior distribution of the vector of parameters of the model, i.e. 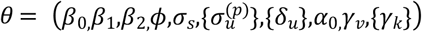. We fitted our model using MCMC using the package NIMBLE ([65] see supplementary material for technical details) for the R software, using 4 chains of 50 000 iterations after a burn-in period of 1000 samples. To save some memory space, we thinned the chains by selecting one sample every 20 iterations. We checked the mixing properties of the chains both visually and using the diagnostic of [67]. We checked the goodness of fit by using the approach recommended by Gelman and Meng [68]. Each MCMC iteration generated a sampled value *θ*^*(r)*^ of the parameter vector *θ*. For each simulated value *θ*^*(r)*^, we simulated a virtual dataset and calculated a summary statistic on it; we then compared the observed statistics with the distribution of simulated values. We used several summary statistics: the number of fawns in each of the three tick-burden classes in total, for each possible age, and in each year. In all cases, the observed values were within the limits of the 95% credible intervals.

## Results

In total, we collected data of tick infestation on 1043 fawns. Age of individuals was estimated between 1 and 20 days (mean ± SD = 4.01 ± 3.05; min = 1, max = 20) during the 1992-2018 period. Only fawns aged between 1 and 8 days were kept for our analysis, as older fawns were rare in our data (8% of the total dataset). For the following analyses, we only used data from the 965 fawns less than 8 days old (mean = 3.35 +/-2.02) when captured (Fig 3). Among them, 696 fawns had <10 ticks (26 fawns/year in average, SD = 11), 167 had 10 to 19 ticks (7 fawns/year, SD = 3) and 102 had ≥20 ticks (4 fawns/year, SD = 3).

**Fig 3.**
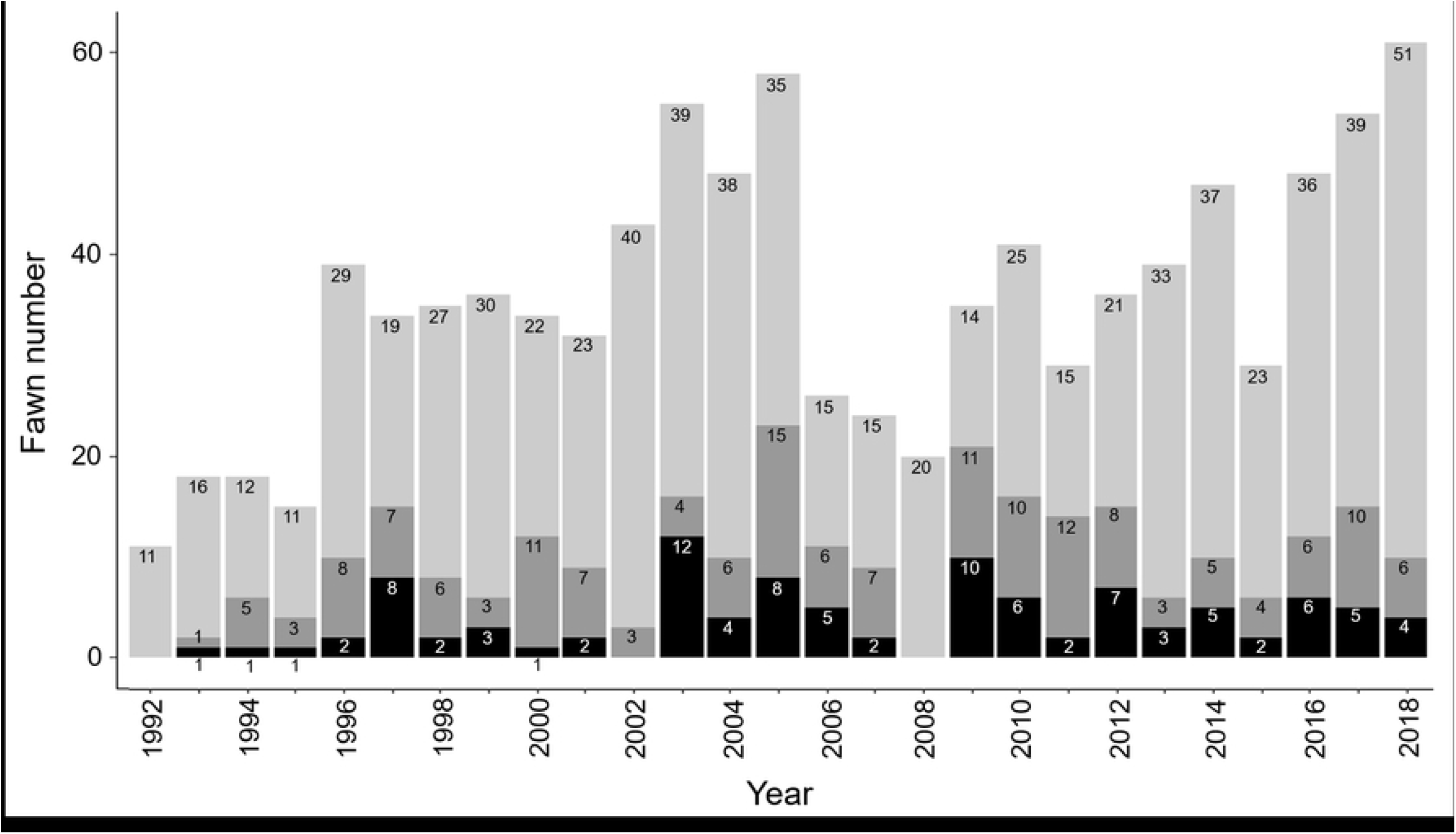
Number of fawns studied per year in the Trois-Fontaines Forest between 1992 and 2018. The three gray scales correspond to the three tick burden classes (gray light = less than 10 ticks; dark gray = between 10 and 20 ticks; black = more than 20 ticks).

The model including an effect of the variables « >5 days old », « humidity » and random effect of the « year » was considered as the best model for the infestation rate of fawns. We did not highlight any effect of the temperature (coefficient = 0.00 [-0.04; 0.04]_95%CI_) and of the roe deer density (coefficient = 0.002 [-0.001; 0.005]_95%CI_) on the instantaneous infestation rate per unit of body surface area. Moreover, all the coefficients associated to habitat types were characterized by 95% CI including 0 – they were all characterized by a point estimate comprised between -0.1 and 0.1 and by a standard error equal to 0.11, see supplementary material). All the estimated parameters of the final model are presented in table 1.

**Table 1.** Estimated top parameters for the final model of the infestation of fawns by ticks in the Trois-Fontaines Forest. For each parameter, we give the notation used in the text, a short description, the point estimate (corresponding to the mean of the posterior distribution) and the 95% credible interval.

| Description | Notation | Est. | 95% CI |
| --- | --- | --- | --- |
| **Intercept in infestation rate log-linear model** | $\alpha_0$ | 2.853 | 2.68 ; 3.02 |
| **Parameters of the growth model** | $\beta_0$ | 0.176 | 0.174 ; 0.178 |
| | $\beta_1$ | 0.014 | 0.013 ; 0.014 |
| | $\beta_2$ | -0.001 | -0.001 ; -0.001 |
| | $\sigma_s$ | 0.021 | 0.02 ; 0.022 |
| **Effect of >5 days old** | $\gamma_v$ | -17.7 | -46.6 ; -2.6 |
| **Dispersion parameter of individuals random effects on the infection rate** | $\phi$ | 0.68 | 0.53 ; 0.85 |
| Effect of humidity in infestation rate log-linear model | $\gamma_k$ | 0.01 | -0.006 ; 0.025 |
| Standard deviation $\sigma_u^{(p)}$ of the years random effects in infestation rate log-linear model for each period p | $\sigma_u^{(1992-1999)}$ | 0.438 | 0.044 ; 1.141 |
| | $\sigma_u^{(2000-2009)}$ | 0.822 | 0.422 ; 1.543 |
| | $\sigma_u^{(2010-2018)}$ | 0.165 | 0.011 ; 0.462 |

During the first few days of life, fawns acquire ticks as they grow. Note that the coefficient *γ*_*v*_, characterizing the rate of infestation for old fawns, was particularly low: this indicates that the instantaneous risk of infestation becomes equal to zero after the first 5 days of life (Fig 4).

**Fig 4.**
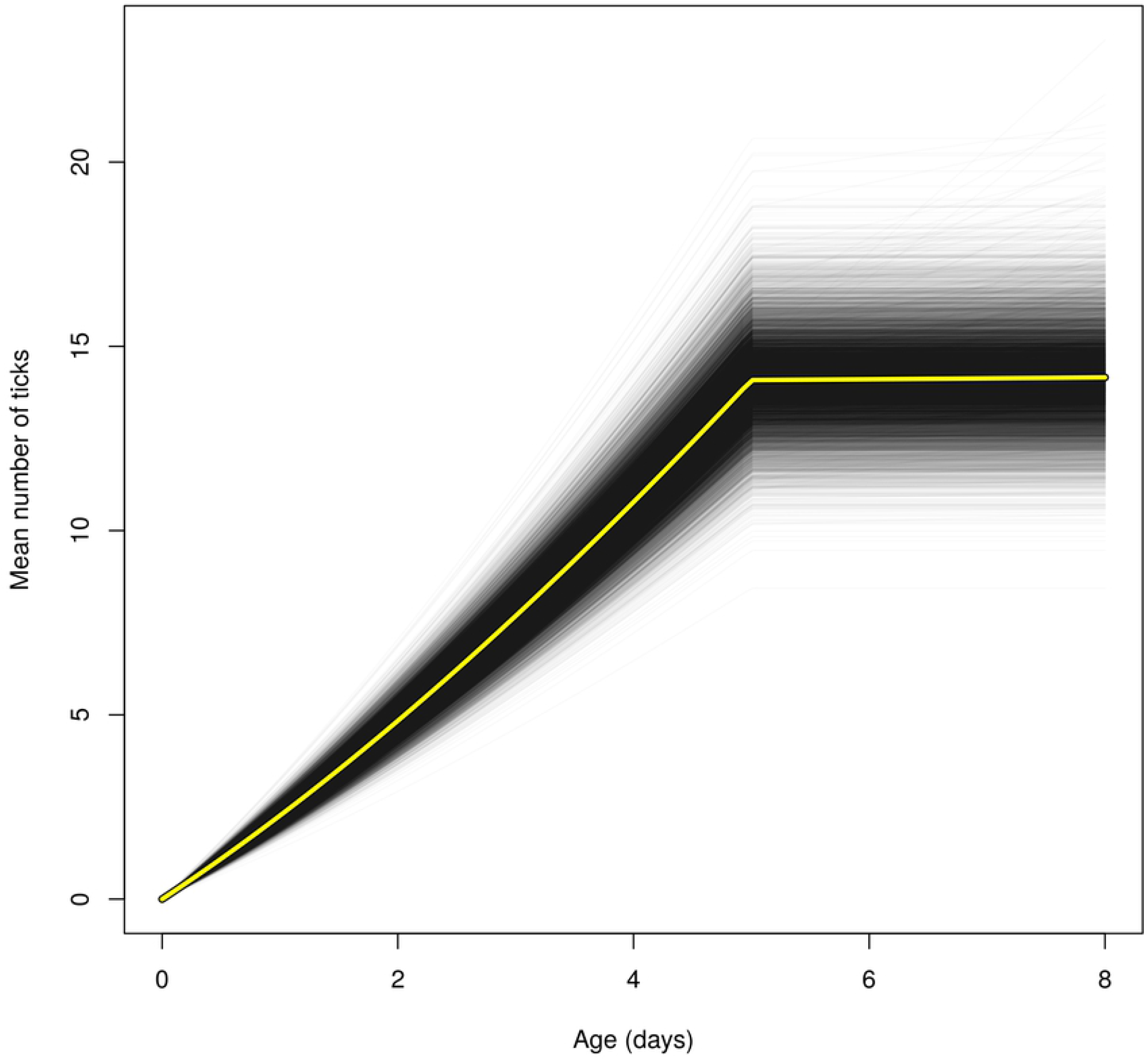
Mean number of ticks expected under our final model (see text) on an average sized fawn of a given age in the Trois-Fontaines Forest during an “average” year (i.e. a fictious year characterized by a random effect equal to 0).

Each thin black curve corresponds to the expected number of ticks obtained for one MCMC iteration (we have 10000 iterations displayed on this plot). The yellow curve corresponds to the mean curve.

The effect of humidity seemed only marginally significant (coefficient = 0.010 [-0.006; 0.025] _95%CI_). When we compared the model including the effect of humidity with a model excluding it with the Watanabe-Akaike Information Criterion (WAIC; [67]), we could not detect any significant difference between the two models (model with humidity: WAIC = 1373.8, SE = 1.57; model without humidity: WAIC = 1374.6, SE = 1.57), so that the two models were equally likely according to this criterion. We therefore decided to keep humidity in the model.

The mean infestation rate was highly variable between years during the 10 years following the hurricane Lothar. Note that this between-year variation was particularly low from 2010 to 2018, the period prior to the hurricane Lothar being intermediate (Fig 5).

**Fig 5.**
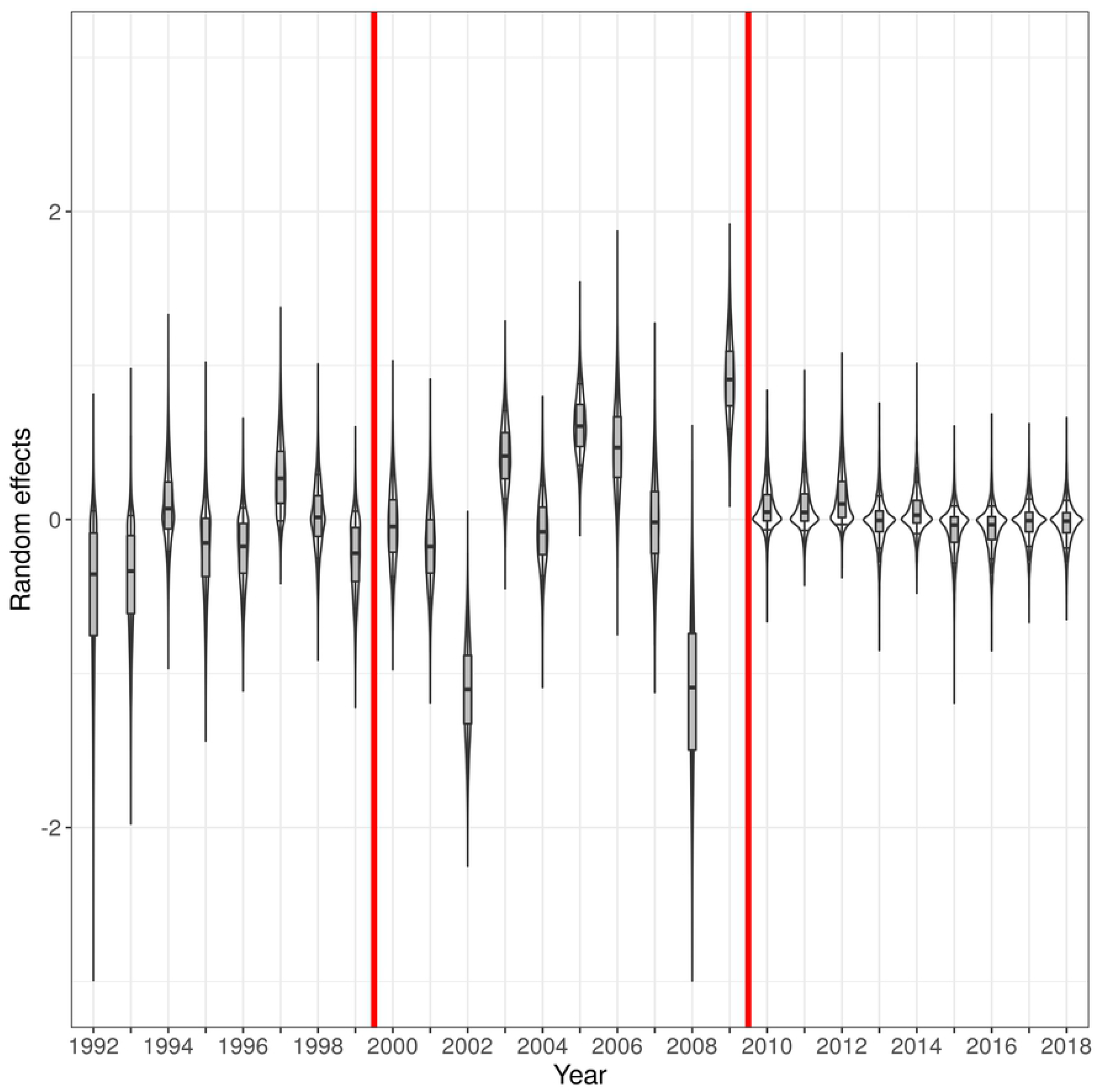
Posterior distribution of the (random) effects *δ*_*u*(j)_ of the year on the instantaneous infestation rate per unit of body size area of fawns by ticks in the Trois-Fontaines Forest.

The vertical red lines delimit the three periods: (i) period 1 corresponded to the period before hurricane Lothar (between 1992–1999), (ii) period 2 corresponded to the 10 years following this hurricane (between 2000 and 2009), (iii) period 3 corresponded to the later years (2010 to 2018). For a given year, the posterior distribution is represented by a boxplot superposed to a violin plot (i.e. for a given random effect, a kernel smoothing of the MCMC simulations of a random year effect rotated and put on both sides of a vertical line, mirroring each other).

A substantial amount of variation in our data remain unexplained. Indeed, the coefficient *φ* characterizing the Gamma distribution of the individual random effects was lower than 1. The parameter *φ* is inversely related to the variance of the Gamma distribution (with a mean = 1 and a variance = 1/φ). A small value of *φ* indicates that the unexplained between-individual random differences in sensibilities are large. And indeed, the Spearman correlation coefficient between the observed dataset and the datasets simulated under our model to assess the goodness of fit of the model was rather small (*ρ* = 0.17 [0.1; 0.24] _95%CI_), confirming that there was still a lot of unexplained variability in our data.

## Discussion

The number of ticks on fawns below the age of 8 days was frequently low. Only 28% of total observations (n = 965) had 10 ticks and more, and 72% of observations had less than 9 ticks (Fig 3). The hiding strategy of roe deer fawns in their first days of life may be effective towards avoiding parasites such as ticks. Indeed, to increase their chance of survival, they stay hidden in the vegetation to minimize the risk of predation [49]. As a result, they have low mobility in the first days of life, leading to a low probability to touch the vegetation where ticks are questing. From the 6^th^ day of life, our study suggests that the level of tick burden becomes constant (Fig 4). This could be explained by the depletion of questing ticks in the habitat of fawns and the turnover of ticks feeding on fawns. Indeed, as ticks take their blood meal on hosts for few days (i.e., in average 3 days for nymphs and larvae and 7 days for adults; [16]) before falling to the ground, where they molt or lay eggs [15], the number of ticks gained after this age is probably counterbalanced by the number of ticks that detach themselves and fall off.

Factors linked to environmental conditions therefore seem to partly drive the between-individual heterogeneity of infestation by ticks. As expected, the humidity rate was positively related to the tick number on fawns. Based on our model, a 20% increase of humidity leads to an increase of 1.2 ticks on fawns in average (i.e., a humidity coefficient equal to 0.01 which is equivalent to a mean number of ticks multiplied by exp(0.01 × 20) = 1.2). This result is in agreement with the positive influence of the relative humidity on *I. ricinus* survival and activity, which can lead to a greater presence of questing ticks in the environment [21,22]. Ticks need at least 70-80% humidity to survive off-host [7]. In this study, the humidity effect was only marginally significant. Our model showed that a 20% increase in humidity (which is large, since humidity varied between 57% and 90% in our area) leads to an increase of 20% of the number of ticks. Since there was in average between 10 and 15 ticks per animal (Fig 4), a 20% increase in the number of ticks corresponds to only 2 to 3 more ticks in average. This may explain our difficulty to identify clearly this effect with our three broad classes of tick-burden.

Tick development and questing activity are strongly temperature dependent [17,24]. Ticks do not tolerate dry and hot conditions due to the risk of desiccation, and below 1°C, they are not active and most of the time goes into wintering [22,24,68]. Thus, climatic conditions are decisive for the survival, development and activity of *Ixodes ricinus*, and this depends on geographic locations and habitat characteristics. We therefore expected to detect an effect of the temperature on the infestation rate. However, no relationship was observed, which could be explained by the lack of power caused by the absence of a precise measure of the number of ticks on fawns.

Similarly, we were not able to identify any effect of the habitat structure on the infestation rate of fawns in our study, and could not validate our second hypothesis. However, it is known that habitat structure can impact tick abundance and distribution, particularly in forest habitat [17]. Indeed, forest vegetation provides a more stable microclimate compared to open habitats, with less extreme variation in climate and mortality risk for ticks [69]. Ground vegetation retains moisture and provides shade, which are very important for tick survival and their water balance. It explains why ticks are mainly found in leaf litter and low layer of vegetation in forests [7,25,43]. Indeed, a forest with important overstory and shrub cover creates a favorable microclimate, and a great depth of litter, and constitutes ideal conditions for their survival [29]. In this study, our inability to isolate any effect of the habitat structure is probably due to the inappropriate scale of habitat measurement: we mapped the vegetation at the resolution of the forest plot, which is probably too large to identify such an effect. Actually, the characteristics at the microhabitat level – i.e., whether the vegetation structure within a few meters around the fawn provides the fawn a thermal protection [72] and low light penetration [73] -- would be more accurate to see differences of infestation rate in terms of vegetation structure. Thus, to better understand how weather factors affect the infestation rate of fawns in a forest habitat, we should look at the tick during their free stages of life in our study area, at a finer spatial scale and consider microclimate conditions.

The level of tick burden on fawns varied according to the decade of our study (Fig 5). The average infestation rate thus exhibits low inter-annual variability during the periods 1992-1999 and 2011-2018. However, there is a stronger interannual variability over the period 2000-2010. This could be linked to the hurricane Lothar, which took place in December 1999. This event has considerably changed the forest structure, by creating numerous openings patches, modifying forest dynamics and habitat use by roe deer [60]. Prior to the hurricane, the Trois-Fontaines forest appeared homogeneous with no strongly different habitat types. After the hurricane, microhabitats and microclimatic conditions became more spatially heterogeneous [71]. Thus, fawns captured between 2000 and 2010 were probably found in habitats with very different characteristics, which influenced the estimated infestation rate of individuals by ticks. Knowing that deer are hosts for both adult and nymph ticks, we first thought that this high interannual variability in infestation rate could have been caused by a variation in roe deer density [72]. Since deer, mainly red deer and roe deer, are the main host of adult *I. ricinus* ticks, their density and distribution can be strong drivers of the location of ticks in the environment [10]. Large-scale exclusion of deer dramatically reduces the abundance of infectious stage of ticks in the environment [73]. However, contrary to our initial hypothesis, we did not find a link between this factor and the level of tick infestation. As stated by Carpi *et al*. [77], the tick infestation of roe deer is not necessarily dependent of the roe deer density. Tick populations can be maintained with a small density of deer, and ticks in the environment did not automatically increase with the abundance of deer [44,75]. Furthermore, during its life cycle, *Ixodes ricinus* can parasitize a large number of vertebrates including mammals (e.g., fox, rabbit, squirrel, etc.) or even birds (e.g., blackbird, pheasant, harrier, etc.) [30]. Thus, to better understand the effect of host density, we should in theory consider the density of all possible hosts of this tick species, which is difficult to monitor.

In summary, the level of tick burden became constant when fawns were 5 days old and more. Humidity was significant but no effects of temperature, vegetation structure or roe deer density were found on the tick burden of fawns. However, we noticed a strong heterogeneity of tick burden between years suggesting other variables could be involved, such as local density of various host species or habitat characteristics of fawns (i.e., bedding sites), which may be important for tick activity. Studies at finer scales need to be carried out to understand why some fawns are more parasitized than others. This would require additional data on ticks in the environment, at different times of the year, which would allow a more precise study to improve our understanding of tick infection in ungulate newborns.

## Acknowledgments

We are grateful to the volunteers and technicians for participating in the fawn captures and data collection at TEE of Trois-Fontaines (France). We also thank Jean-Michel Gaillard for the roe deer density data at Trois-Fontaines. This study was conducted in the context of a PhD co-funded by the OFB and the VetAgro Sup under partnership agreement.

